# Lattice arrangement of myosin filaments correlates with fiber type in rat skeletal muscle

**DOI:** 10.1101/720300

**Authors:** Weikang Ma, Kyoung Hwan Lee, Shixin Yang, Thomas Irving, Roger Craig

## Abstract

The thick (myosin-containing) filaments of vertebrate skeletal muscle are arranged in a hexagonal lattice, interleaved with an array of thin (actin-containing) filaments with which they interact to produce contraction. X-ray diffraction and electron microscopy have shown that there are two types of thick filament lattice. In the simple lattice, all filaments have the same orientation about their long axis, while in the super lattice, nearest neighbors have rotations differing by 60°. Tetrapods (amphibians, reptiles, birds, mammals) typically have only a super lattice, while the simple lattice is confined to fish. We have carried out X-ray diffraction and electron microscopy of the soleus (SOL) and extensor digitorum longus (EDL) muscles of the rat and found that while the EDL has a super-lattice, as expected, the SOL has a simple lattice. The EDL and SOL of the rat are unusual in being essentially pure fast and slow muscles respectively. The mixed fiber content of most tetrapod muscles and/or lattice disorder may explain why the simple lattice has not been apparent in these vertebrates before. This is supported by only weak simple lattice diffraction in the X-ray pattern of mouse SOL, which has a greater mix of fiber types than rat. We conclude that the simple lattice might be common in tetrapods. The correlation between fiber type and filament lattice arrangement suggests that the lattice arrangement may contribute to the functional properties of a muscle.

**Summary:** The three-dimensional arrangement of thick filaments in skeletal muscle is studied by X-ray diffraction and electron microscopy. A correlation is found between thick filament lattice type (simple or super lattice) and fiber type (fast/slow). This suggests that lattice organization contributes to muscle functional properties

## Introduction

The thick and thin filaments of vertebrate striated muscle are arranged in a double hexagonal lattice, in which each thin filament lies at the trigonal point between three thick filaments [5]. Interaction between myosin heads on the thick filaments and actin subunits of the thin filaments is responsible for the relative filament sliding that generates contraction [6]. Electron microscopy (EM) combined with X-ray diffraction has shown that the thick filaments are organized in one of two ways [1, 3, 7, 8]. In one, all filaments have the same rotational orientation (a “simple” lattice), while in the other, nearest neighbors have orientations differing by 60°, and only next-nearest neighbors have equivalent orientations (a “super” lattice). These different lattices are recognized in the electron microscope by the orientation of thick filament triangular profiles seen in transverse sections of the bare zone – the region of the thick filament next to the M-line, lacking myosin heads (Fig. 1A) [3, 7, 8]. The lattices can also be distinguished in X-ray diffraction patterns, where myosin layer lines, arising from pseudo-helical organization of the myosin heads [1], are sampled either at the same radial positions as the equatorial reflections (simple lattice) or in a more complex pattern (super lattice; Fig. 1B; [1, 8]). EM analysis has revealed a simple rule for filament orientations in the super lattice: for any group of three nearest neighbor filaments, in a line or in a triangle, two have the same orientation, while the third is rotated 60º (the no-three-alike rule) [3, 7, 8].

**Figure 1.**
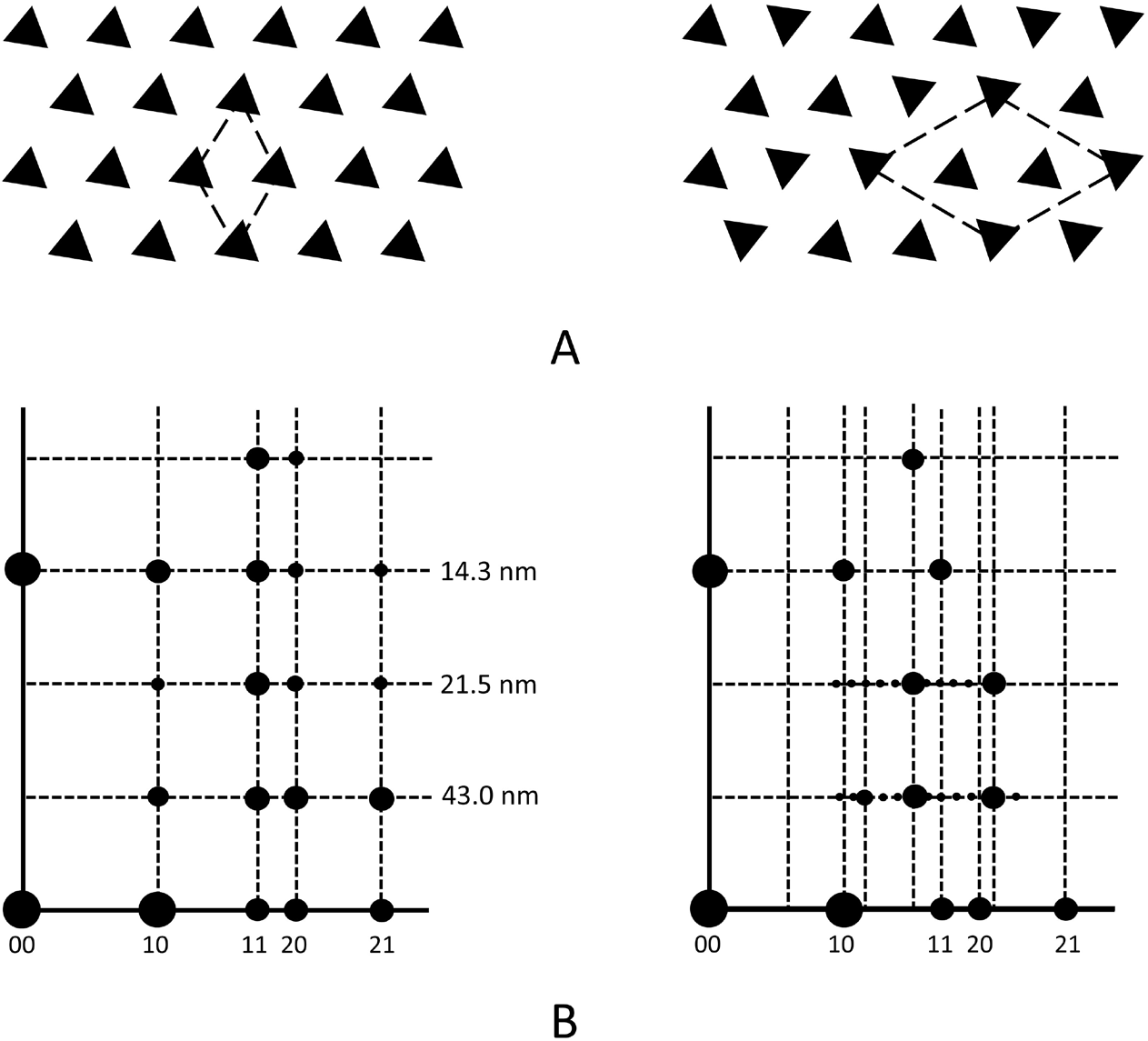
Simple and super lattice models (left and right, respectively). **A.** Models of transverse sections of thick filament bare zones in electron micrographs. **B.** Sampling of intensity on myosin layer lines of X-ray diffraction pattern. 10, 11, etc. show positions of reflections on equator. 43.0, 21.5, 14.3 nm show positions of 1^st^, 2^nd^ and 3^rd^ myosin layer lines. In simple lattice, note alignment of layer line sampled spots with corresponding equatorial reflections; in the case of the super lattice, the sampling is more complex. Based on [3] and [4], with permission.

The super lattice arrangement was first recognized in X-ray diffraction patterns of frog skeletal (sartorius) muscle [1] and was confirmed in electron micrographs of the same muscle [7]. Other tetrapods (amphibians, reptiles, birds, mammals) examined since then also typically exhibit only a super lattice [3]. The super lattice is generally not very well ordered and extends over only a small number of unit cells [7]. The simple lattice is observed specifically in fish, particularly teleosts (the predominant group of bony fish) [3, 8, 9]. However, some primitive fish (e.g. hagfish, lampreys, sharks and rays) have been shown to have a super lattice, suggesting that this form evolved earlier [3, 8]. Interestingly, sharks and rays exhibit simple and super lattices in the same animal, where they correlate with muscle type, the simple lattice being present in red (slow) fibers, and the super lattice in the white (fast) fibers [3, 8].

Here we study the thick filament lattice arrangement in a mammal (rat) and determine whether this differs between their fast and slow muscles. Most mammals have a mixture of fast and slow fiber types in individual muscles, complicating such analysis. The rat is unusual in that its soleus (SOL) and extensor digitorum longus (EDL) muscles have a relatively homogeneous population of fiber types. The SOL contains 85-95% slow type 1 fibers, with 5-15% fast IIA [10, 11], while the EDL contains ~100% fast fibers (Type IIA, IIB and IID/X) [11, 12]. Thus it is possible to get relatively pure (i.e. fast or slow muscle) X-ray diffraction patterns and EM images from entire anatomical muscles. The mouse, in contrast, is ~100% fast in EDL (type IIB/DB/AD,X) [13, 14], but in the SOL it is ~35-40% slow (Type I) and 55-65% fast IIA/D/B/X [13–15]. Our results from rat muscle show that slow fibers have a simple lattice, while the fast fibers have a (relatively poorly ordered) super lattice. This correlation suggests that the specific rotational arrangement of myosin filaments may be one of the determinants of muscle fiber functional properties (e.g. level of tension) in mammals [3, 8].

## Materials and Methods

### Muscle preparation

Rats (Sprague-Dawley, 150 g) and mice (C57BL/6J, 25g) were euthanized by CO_2_ asphyxiation followed by cervical dislocation according to IACUC-approved protocols of the University of Massachusetts Medical School and Illinois Institute of Technology. The skin was removed and the hind limbs were separated. The hind limbs were placed in a dish with Ringer solution (145 mM NaCl, 2.5 mM KCl, 1.0 mM MgSO_4_, 1.0 mM CaCl_2_, 10.0 mM HEPES, 11 mM glucose (pH 7.4) and perfused with 100% oxygen. For intact muscle experiments, EDL and SOL muscles were rapidly dissected, and tied with one suture on each end of the muscle. For skinned muscle experiments, EDL and SOL muscles were pinned to a Sylgard^TM^ substrate at approximately physiological length during skinning overnight at 4° C with gentle agitation in skinning solution (40 mM BES, 10 mM EGTA, 6.56 mM MgCl_2_, 5.88 mM Na-ATP, 46.35 mM K-propionate, 15 mM creatine phosphate and 1% Triton X-100). The muscle was rinsed in relaxing solution (skinning solution lacking detergent) before being placed on the X-ray diffraction apparatus.

### X-ray diffraction

Intact or skinned muscles were placed in a specimen chamber containing Ringer’s or relaxing solution and exposed to a collimated X-ray beam of wavelength 0.1033 nm at the BioCAT beam line at the Advanced Photon Source, Argonne National Laboratory [16]. One suture was attached to a hook inside the chamber, and the other to a dual-model motor/force transducer level (Aurora Scientific Inc, Model 300_LR). The muscles were adjusted to optimal length (the length for generating maximal force) by an X-Y-Z positioner attached to the motor/force transducer. For intact muscle experiments, the experimental chamber was oxygenated throughout the experiment. The rat muscle diffraction patterns were recorded on a Mar 165 CCD detector (Rayonix), and the mouse diffraction patterns on a Pilatus 3 1M detector (Dectris, Inc.). The diffraction patterns were quadrant-folded and background subtracted using the MuscleX software package developed at BioCAT [17]. The contrast and brightness of the patterns were adjusted in ImageJ to optimally reveal features of interest. The strong equatorial reflections are shown on a reduced scale for clarity.

### Electron microscopy

Live muscles tied to sticks were fixed for 1 hr at 4º C in 3% paraformaldehyde/ 0.1 % glutaraldehyde followed by 2.5% glutaraldehyde (both in 0.1M phosphate buffer, pH 7.4), post-fixed in 1% OsO_4_ in 0.1 M sodium cacodylate, dehydrated in an ethanol series and embedded in Epon. Transverse sections 65 nm thick were cut on a diamond knife using a Leica UC7 ultramicrotome. Sections were stained with uranyl acetate and lead citrate, and examined at 120 kV in a Tecnai Spirit transmission electron microscope. Images from myofibrils sectioned in the bare zone region of the thick filament (Fig. S1) were recorded with a pixel size of 0.41 nm on a Gatan Erlangshen CCD camera.

### Image analysis

To determine thick filament rotational orientation objectively, an approach was developed based on single particle analysis methods currently widely used in cryo-EM studies [18]. Each triangular filament profile in the transverse image of a bare zone was treated as a single particle. Particles were manually selected and boxed, subjected to 2D classification, then a class average was computed for each class, all using RELION [19]. SPIDER was used for subsequent image processing [20]. The best 2D class average for each micrograph was then used as a reference for projection matching to all the filaments in an image. The rotational angle of a filament that best matched the reference was determined by cross-correlation. This angle defined the rotational orientation of each filament. A triangle was then superposed on each filament in the original image with the determined orientation (Fig. 3C, E). Finally, the superposed triangles representing each filament were transferred to an empty background to clearly reveal each filament orientation (Fig. 3D, F).

To determine the lattice type, a statistical analysis of the orientations of the triangles in each image was carried out. The relative orientations of the triangles in any hexagon of the lattice were determined by selecting one triangle as the central filament, and calculating the distance between this triangle and the others in the same image. The nearest neighbor surrounding filaments were selected as those within a threshold distance of 40 nm (slightly greater than the center-to-center distance of two adjacent triangles), ensuring that only nearest neighbor filaments of a hexagon were selected. The average deviation angle between the central filament and its six surrounding neighbors (green circles, Fig. 3D, F) was calculated using the azimuthal angle from the orientation determination above. The sign of the deviation angle was not taken into account. This step was repeated for each triangle in the image.

In the ideal simple lattice, all triangles have the same orientation (Fig. 1). This was approximated by filaments in the rat SOL, which showed only small deviations from the reference (Fig. 3D). In the super lattice, adjacent triangles have either a 0º or 60º difference in angle (Fig. 1). Therefore, two statistical distributions can be calculated. One is the average deviation for all six filaments surrounding a central filament, as for the simple lattice above. For the EDL, the peak in the distribution histogram was close to 30º (Fig. 4D). The other distribution is the pairwise deviation between one filament and the adjacent filament. To automatically select filaments for this measurement, the distance between the one filament and its neighbor had to be less than the threshold of 40 nm. The deviation between filaments of a pair was then calculated using the orientation angle for the two filaments. The histogram shows one peak close to 0º and another close to 60º for rat EDL (Fig. 4E).

## Results

### X-ray diffraction

Axial X-ray diffraction patterns were recorded from both intact and freshly skinned rat SOL and EDL muscles under relaxing conditions. Results were similar for both (Figs. 2, S2), and we describe only intact muscle below. The equator of the patterns showed the typical strong 10 and weaker 11 reflections of relaxed muscle, consistent with the close association of myosin heads with the thick filament backbone, with minimal interaction with actin [5] (Fig. 2). At higher exposure, weaker reflections (e.g. 20, 21; Fig. 1B) were observed further out on the equator (not visible when scale adjusted for clear 1,0 and 1,1 reflections). The positions of these equatorial reflections reflect the hexagonal arrangement and spacing of myosin filaments, with interleaving actin filaments, but do not provide information on the relative orientations of the myosin filaments. In addition to the equator, the patterns also showed a series of layer-lines (ML1-ML6) indexing on a repeat of 430 Å, reflecting the pseudo-helical organization of the myosin heads on the thick filaments (Fig. 2A, B cf. Fig. 1B; [1]). The distribution of intensity along the layer lines was quite different between SOL and EDL. In SOL, lattice sampling produced intensity that was strongest at radial positions corresponding to those of the equatorial reflections (Fig. 2A). This was most obvious on the first layer line, but also evident on higher order layer lines (Fig. 2A, rectangular boxes; Fig. S3). This lattice sampling appeared similar to that in the simple lattice X-ray patterns from fish muscle [8, 21]. In the EDL, the distribution of intensity on the first layer line was relatively continuous, with little sign of lattice sampling, suggesting a rotationally less coherent lattice of thick filaments (Fig. 2B). Careful inspection, however, revealed weak lattice spots on ML1 that did not align with the equatorial reflections, and looked similar to the pattern first described for frog skeletal muscle, which has a super lattice (Fig. S4) [1, 8]. Intensity distribution was different on the higher layer lines. We conclude that the SOL and EDL have simple and super lattice thick filament arrangements, respectively, and that these may be due to the different fiber types (slow and fast) that predominate in these muscles in the rat. The weakness of the superlattice reflections on ML1 in the EDL pattern suggests that the super lattice is significantly disordered [7, 8].

**Figure 2.**
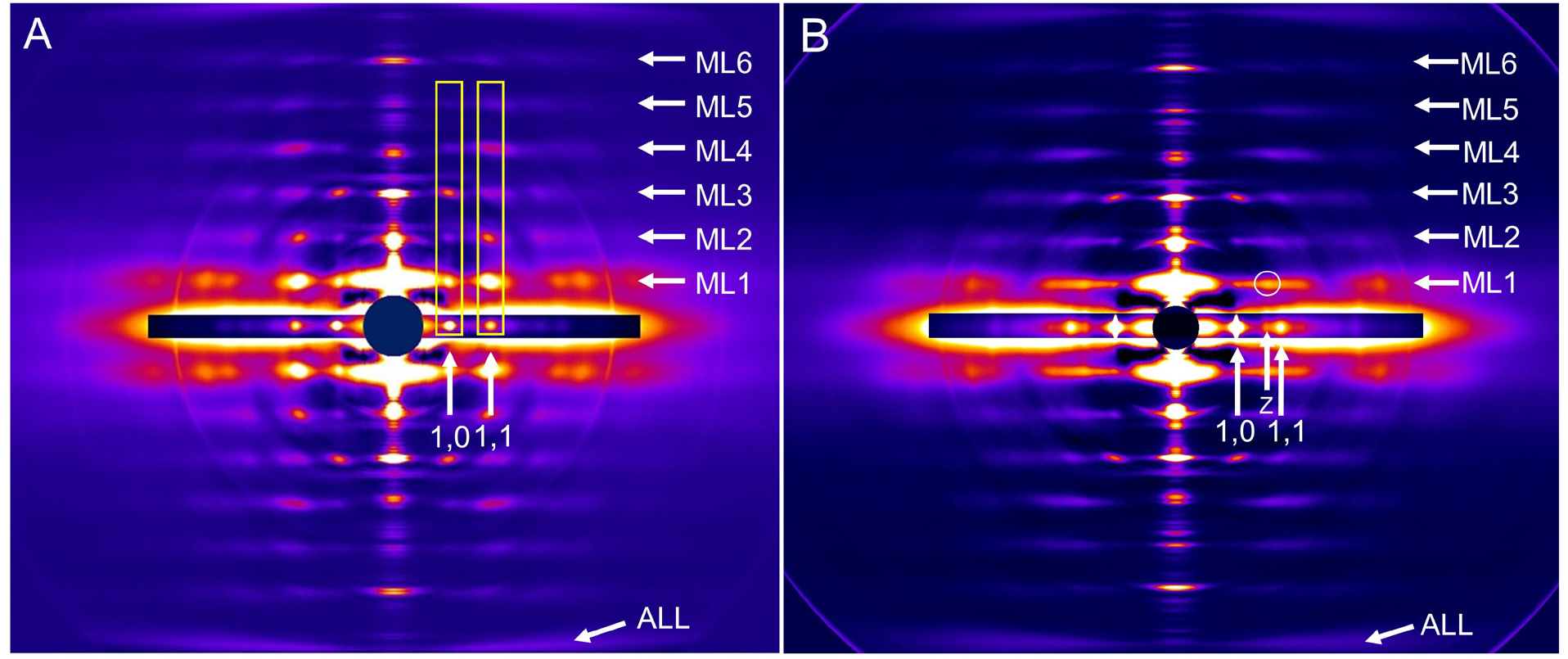
X-ray diffraction patterns of rat SOL (A) and EDL (B) muscles. ML1-6 indicate positions of myosin layer lines (repeat 43 nm), and ALL, actin 5.9 nm layer line. Horizontal inserts on equator show shorter exposures to reveal the high intensity equatorial 1,0, and 1,1 and Z-line reflections. Prominent lattice sampling is seen in SOL at the same positions as equatorial reflections (vertical rectangles aligning with 1,0 and 1,1 reflections). This indicates a simple lattice of myosin filaments. Lattice sampling is largely absent in EDL, except for weak super-lattice spot circled. The pattern suggests a local super-lattice with disorder.

### Electron microscopy

Rat SOL and EDL muscles were fixed, embedded and transversely sectioned for examination by transmission EM. Sections were studied in the “bare zone” of the thick filaments, just to each side of the M-line (Fig. S1). The bare zone lacks both myosin heads and the M-line bridges that link thick filaments to each other. It therefore provides the clearest visualization of filament rotational orientation, recognized by the triangular profiles of the thick filament backbones that reflect their 3-fold symmetry (Figs. 1A, 3A, B; [7]). The relative orientations of these profiles provides a direct visualization of the type of filament lattice (Fig. 1A). However, even with the thin (65 nm) sections that we used, some portion of the M-line bridges and/or myosin heads will be included, as the bare zone region on each side of the M-line is only about 30 nm long. The M-line and/or myosin heads presumably account for the additional, low-density material surrounding the thick filaments, which can diminish the clarity of the triangular profiles (Fig. 3A, B).

**Figure 3.**
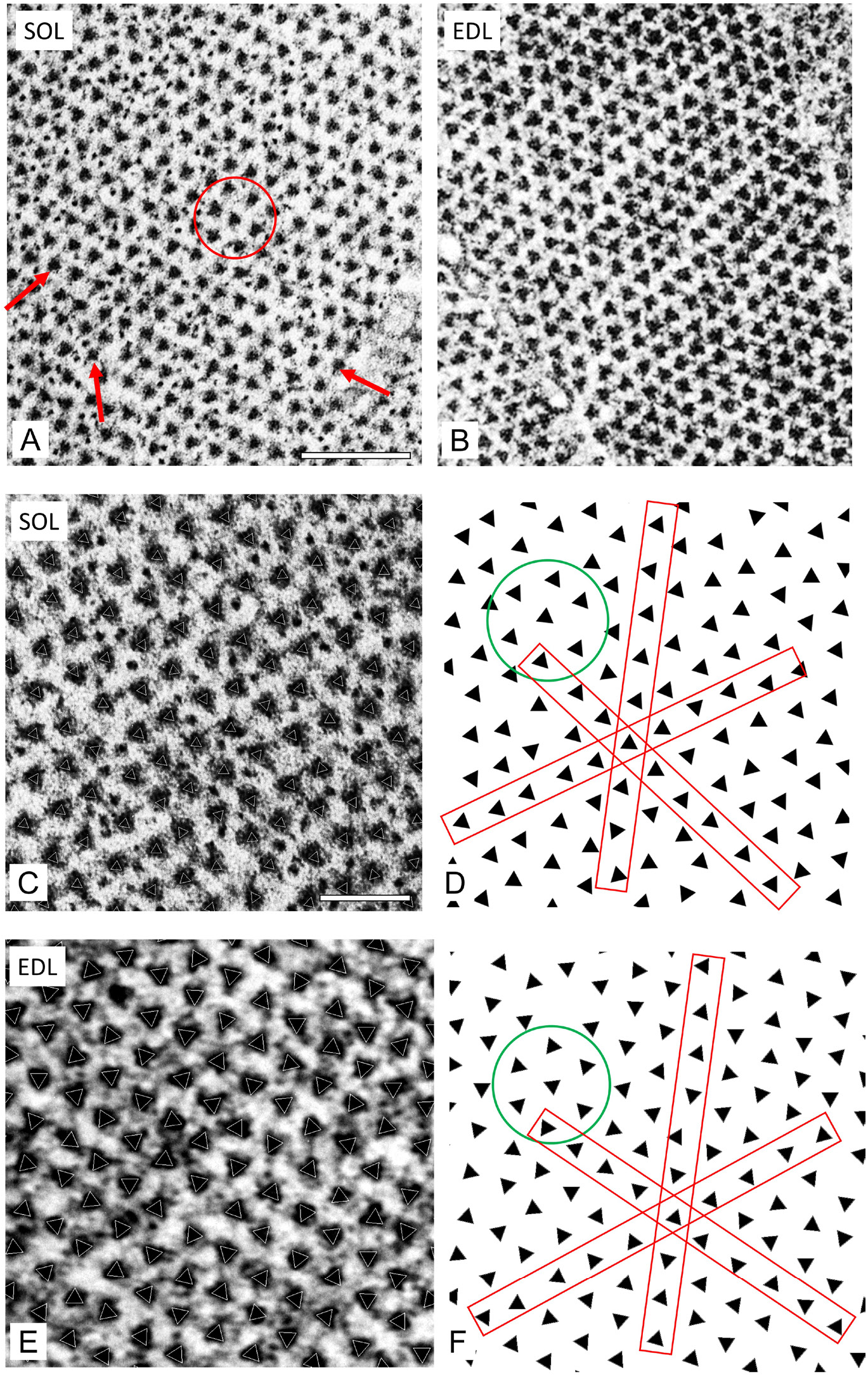
Transverse sections of bare zone of rat SOL and EDL revealing lattice type. **A.** SOL shows triangular filament profiles pointing in approximately the same direction along all three lattice planes (arrows) and within a hexagon (circle). This suggests a simple lattice. **B.** EDL shows triangular profiles with variable orientations, which are analyzed below. **C-F.** SOL and EDL sections at higher magnification analyzed by projection matching. **C, E.** Raw images with superposed white triangles representing orientation determined by projection matching (see text). **D, F.** Triangles extracted from C, **E.** reveal orientations most clearly. **D.** shows triangles with similar orientations in all 3 planes (red rectangles) and in hexagon (circle). **F.** shows triangles with varying orientations, mostly following the no-3-alike rule. Scale bars, 100 nm in A,B; 200 nm in C-F.

In rat SOL, neighboring thick filaments in well-contrasted bare zones typically had approximately the same rotational orientation (Fig. 3A). This was apparent to the eye by comparing the group of six peripheral and one central filament in a hexagon (Fig. 3A, circle), or by comparing filament orientation along the rows of filaments in any one of the three lattice directions (Fig. 3A, arrows). The filament lattice in EDL had a distinctly different appearance, showing nearest neighbors with varying orientations (Fig. 3B). To provide an objective assessment of filament orientation and to quantify the relative orientations, the rotational angle of each filament was determined computationally using a projection matching approach (see Materials and Methods). A triangle with the orientation thus determined was then superposed on each filament in the image (Figs. 3C, E). To most clearly visualize filament orientations, the triangles were then displayed without the original micrograph (Figs. 3D, F). The results confirmed the visual appearance in Figs. 3A, B. The soleus showed triangles with similar orientations in all 3 planes (red rectangles, Fig. 3D) and within individual hexagons (circle; Fig. 3D), while EDL revealed triangles with varying orientations, which tended to follow the no-3-alike rule (Fig. 3F) [7, 8].

The relative orientations of neighboring filaments were then analyzed statistically by calculating the difference in angle between each filament and its six nearest neighbors and plotting the results as a histogram (Fig. 4). In a simple lattice, all six filaments have the same orientation as the central filament (mean deviation angle 0º; Fig. 4A). The histogram for the SOL micrograph in Fig. 4A showed a peak at about 12º, supporting a simple lattice (expected deviation angle 0º). In a no-3-alike super lattice (Fig. 4C), two of every three adjacent filaments have the same angle and the third is rotated by 60º. The predicted mean deviation angle is therefore 30º. The histogram for the EDL micrograph in Fig. 4D shows a peak close to 30º, consistent with a no-3-alike super lattice in this muscle. However, 30º is also consistent with a random filament orientation. To distinguish between these two possibilities we carried out a second statistical test. In the no-3-alike super lattice, pairs of adjacent filaments should have either the same orientation or differ by 60º, and these should appear in equal numbers. The EDL shows exactly this behavior, supporting a super lattice in this muscle (Fig. 4E).

**Figure 4.**
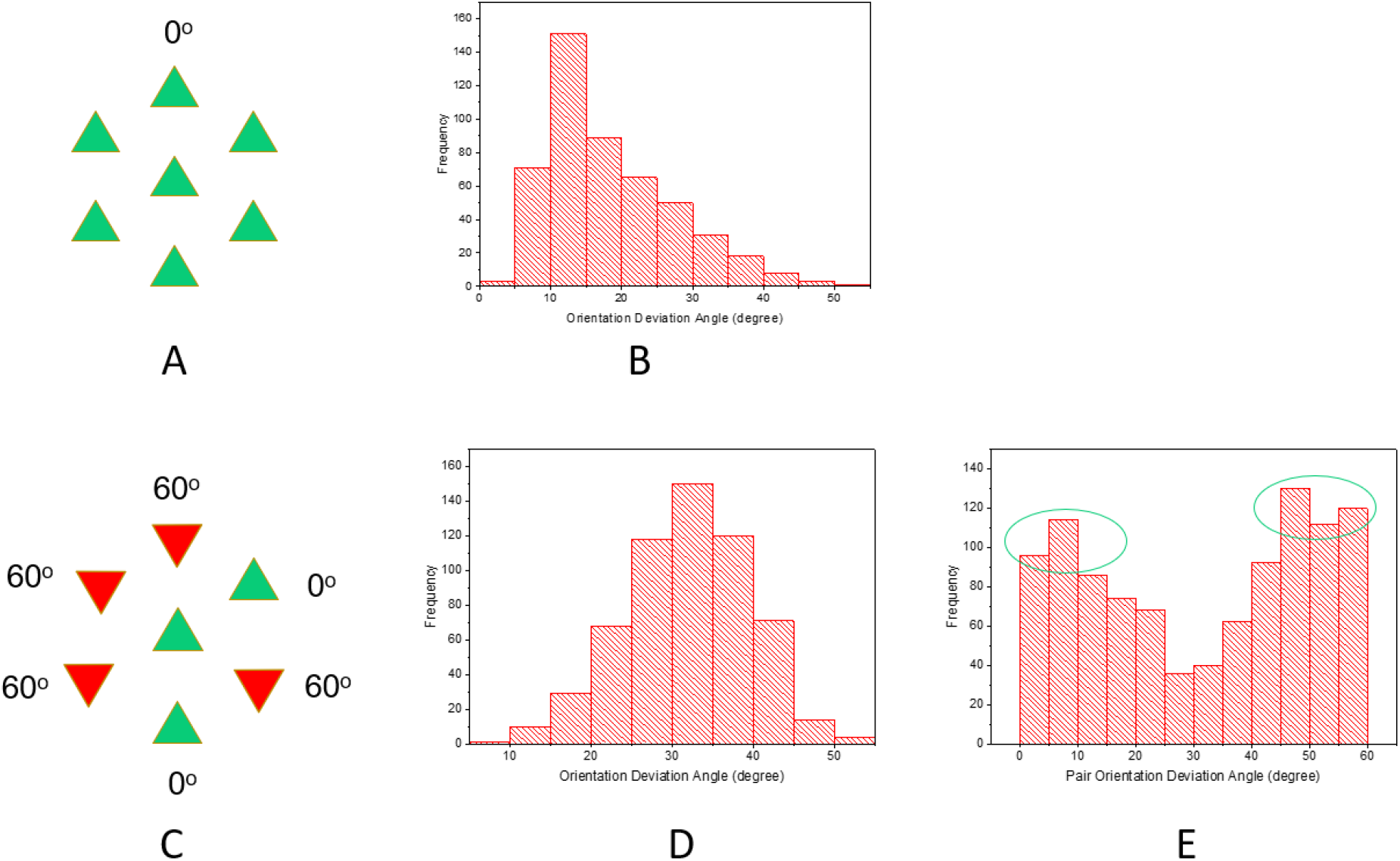
Analysis of filament orientations in one micrograph each of SOL and EDL (see also Fig. S5). **A, C.** Filament orientations in **(A)** simple lattice (all filaments have same orientation; mean deviation angle = 0º) and **(C)** no-3-alike super lattice (in any group of three filaments, in a line or forming a triangle, one filament is rotated by 60º with respect to the other two, which are identical; mean deviation angle for any three adjacent filaments = 30º). **B.** Histogram showing deviation angle for SOL micrograph. The histogram peaks near 10º, close to the expected 0º for a simple lattice. **D.** Deviation angle for three adjacent filaments for EDL micrograph. The peak is close to 30º, as expected for a no-3-alike super lattice. **E.** Deviation angle for adjacent filaments in EDL considered pairwise. There are two peaks, close to 0 and 60º, as expected for a no-3-alike super lattice (see **C.**). Note: the relatively broad spread of the angle data is partly due to imperfections in the filament lattice as seen in bending of the lines of filaments (Fig. 3) and imperfections in particle picking due to thin filament proximity to thick filament. Both factors affected the accuracy of angle determination.

A similar comparison of filament orientations was carried out in a total of 8 micrographs for SOL (3,959 filaments in total) and 8 micrographs for EDL (2,850 filaments). Micrographs were selected for best visibility of bare zone triangular profiles. The aggregate results are shown in Fig. S5, and support the conclusions from the individual micrographs shown in Fig. 4. Thus, electron microscopy supports the simple and super lattices suggested by the X-ray diffraction patterns of the rat SOL and EDL muscles respectively.

## Discussion

Previous EM and X-ray studies showed that vertebrate skeletal muscles have one of two types of thick filament lattice: simple or super. Comparison of different species suggested that tetrapods had only super lattices, and that simple lattices were confined to fish [3, 8]. Our observations show that the soleus muscle of the rat (a tetrapod) also has a simple lattice. They also suggest that this is related to the slow fiber type that predominates in this muscle, as the EDL (a fast muscle) has a super lattice. It will be interesting to see whether other muscles of these rodents also show a correlation between fiber type and lattice type, and whether this association applies to other mammals and to the other tetrapod groups (amphibians, reptiles, birds).

Why has a correlation between fiber type and lattice type not previously been recognized in tetrapods? The main reason is probably the limited number of tetrapod skeletal muscles that have been examined by X-ray diffraction – primarily frog sartorius [1, 22, 23] and rabbit psoas [24] – both fast muscles [25–28]. To our knowledge, only three “pure” tetrapod slow muscles have previously been examined by X-ray diffraction (most muscles are not pure, but have a mixture of fast and slow fibers [29]). A study of rat SOL and EDL, under conditions designed to phosphorylate or dephosphorylate the regulatory light chains, showed little to no lattice sampling on ML1 of SOL, and made no conclusion about the type of myosin filament lattice present [30]; the conditions used for these experiments may have caused significant disordering of the myosin heads, thus weakening the myosin layer lines and their sampling. A comparison of rat SOL and psoas muscles focused on crossbridge behavior and made no mention of lattice structure [31]. Likewise, a comparison of chicken slow (anterior latissimus dorsi – ALD) and fast (posterior latissimus dorsi – PLD) muscles centered on myosin head proximity to thin filaments, but did not study myosin layer lines or comment on the type of filament lattice [32]. Given this paucity of X-ray data on fast and slow muscles, and focus on other aspects of structure when they have been studied, it is possible that a correlation between fiber and lattice type is common but has simply been missed. The relatively pure composition of the rat SOL (~90% slow) and EDL (~100% fast), however, made the lattice/fiber type correlation immediately obvious in our X-ray patterns (Fig. 2).

The importance of a pure fiber composition for detecting lattice type is supported by a previous X-ray study, which was designed to compare the structures of the skeletal muscles of amphibia and mammals [33]. The muscles chosen were frog sartorius and mouse diaphragm. Mouse diaphragm is thought to consist almost entirely (98%) of fast-twitch fatigue-resistant (FFR) type IIA fibers [34]. While frog sartorius showed a super lattice, as previously found [1], the mouse diaphragm showed evidence for a simple lattice—the only other tetrapod muscle reported to have shown this—with lattice sampling similar to that in rat SOL (Fig. 2A) and fish [8]. This finding suggests that the simple lattice may correlate not specifically with slow muscle, as we initially hypothesized based on our rat studies, but with fatigue-resistant fibers, which includes both FFR and slow muscles [29]. Because other mouse muscles were not included in their comparative study [33], it was not recognized that the simple/super lattice dichotomy might relate to fiber type, as our results suggest, rather than animal group (mammal vs. amphibian).

While rat SOL and EDL are typically regarded as slow and fast muscles respectively, there is still some fiber type mixing. Does the interpretation of our X-ray results hold, given this added layer of complexity? Rat SOL and EDL have been analyzed for both myosin heavy chain and fiber type. Labeling with anti-fast and anti-slow myosin antibodies yielded a ratio of 90% slow and 10% fast in the SOL of young rats (~12 weeks) and 95% slow and 5% fast fibers at 1 year old [10]. This is roughly consistent with myosin heavy chain type determined by SDS-PAGE, which yielded 80% type I (slow) fibers and 20% type IIA (“fast”) at ~12 weeks [12]. Similar results are obtained based on myosin ATPase activity and immunocytochemistry [35]. By mass spectrometry, rat SOL (age not reported) had 88% type I fibers and 12% type IIA fibers [11]. Overall these data imply that rat SOL has 80-95% slow, 5-20% fast IIA fibers, and no fast IIB/X/D fibers. Assuming that type I and IIA fibers both have a simple lattice, as discussed above, the lattice sampling from this mixed muscle is consistent with our X-ray data. The rat EDL also has a mixed fiber content: 64% IIB, 25% IIX and 11% IIA (from myosin type by SDS-PAGE; [12]); 76% IIB, 19% IIA, and 5% type I [35]; and 70% IIB, 24% IID/X and 5% IIA by mass spectrometry [11]. IIB/X/D are all considered to be fast fibers [29]. With only ~10% or less IIA fibers, the expected small amount of simple lattice sampling would be difficult to detect in EDL, consistent with our X-ray data. We conclude that our rat diffraction data are consistent with the idea that slow type I and fast type IIA fibers have a simple lattice while the fast fibers (IIB/X/D) have a super lattice.

The dearth of X-ray data on tetrapod lattice types is mirrored by a lack of EM data. To our knowledge, no previous EM study has addressed the thick filament lattices present in the different fiber types of tetrapods, and a simple lattice has not previously been observed by EM in any tetrapod (the study of mouse diaphragm [33] did not use EM and it will be important to confirm the X-ray interpretation of the lattice in this muscle by direct EM observation). It is therefore possible that a simple lattice structure in slow and FFR fibers has simply been overlooked. Early EM studies of fish muscle were similarly limited—to teleosts (the predominant group of bony fish)—and revealed only simple lattices [9]. When comparison was widened to include other, more primitive groups of fish (e.g. hagfish, lampreys, sharks and rays) super lattices were found to be common [3, 8]. A comparable, more detailed survey might show that simple lattices are common in tetrapods.

Our combined EM and X-ray data suggest that the type of thick filament lattice is related to muscle fiber type in the rat. Further work will be necessary to determine whether this correlation extends to other rodents and other tetrapods (including other mammals). We have made a first step to answer this question by studying the SOL and EDL muscles of the mouse. Overall, there is reasonable agreement with the conclusions from the rat (see Supplementary Materials and Figs. S6-S10). However, the results are less definitive due to the mixed fiber content of mouse SOL, the difficulty of identifying different fiber types in the SOL by EM, and apparently greater disorder of the mouse lattices compared with rat. Nevertheless, the mouse data are broadly consistent with the conclusion that low-fatigue fibers (type I and IIA) have a simple lattice while fatigable fibers (type IIB, D, X) have a super lattice. In no case is the lattice perfect, but in both SOL and EDL is substantially disordered (i.e. of limited extent).

Strikingly, sharks and rays in which white and red fibers were compared by EM showed a similar lattice/fiber type correlation [3, 8] to that in the rat. Red fibers (slow) had a simple lattice while white fibers (fast) had a super lattice. This supports the notion that lattice type is connected to fiber type, and suggests that it may occur over a range of vertebrates, including tetrapods as well as fish. However, the correlation is not perfect, as no such connection is found in teleosts, where red and white fibers both have simple lattices [3, 36].

Differences in the physiology of slow and fast fibers have a profound impact on the functioning of skeletal muscle. Slow (type I) fibers play an essential role in postural muscles and muscles that function continuously, without break, for the life of the organism (e.g. respiratory muscles of larger species [29, 37]). They contract more slowly and generate less tension than fast fibers, but they are highly resistant to fatigue [29]. Fast, fatigable fibers (IIB/D/X) are complementary, adding to muscle response when brief but powerful contractions are required [29, 38]. Fast, fatigue-resistant fibers (type IIA) have intermediate properties [29]; although classically described as fast, they are in many ways more akin to slow fibers and quite distinct from IIB/D/X [29]. Differences in mitochondrial content (and the oxidative metabolism that this supports), and in myosin heavy and light chain isoforms (with associated differences in actin-activated myosin ATPase activity) underlie much of this diversity in function. Our results raise the possibility that the structural organization of the myosin filaments might also contribute to the different contractile properties. The proper functioning of relatively slowly contracting/fatigue-resistant muscles (types I, IIA), designed for prolonged activity, may depend in part on high mechanical efficiency of contraction [29]. The arrangement of thick filaments in simple or super lattices will affect the ease with which myosin heads attach to actin filament target zones, which could impact the contractile properties (e.g. speed, tension) of a fiber [3, 8]. Thus the 3D geometry of actin-myosin interactions defined by the simple lattice might favor efficiency, while the super lattice may enable a greater number of actin-myosin interactions, contributing to the greater force production of fast, fatigable fibers [3, 8, 36, 39].

What is responsible for generating the two lattice types found in the different fiber types of rats and mice (and possibly other tetrapods)? Thick filament crosslinking proteins, such as those of the M-line, are likely to be involved, as these provide a means of setting thick filaments with specific orientations relative to each other. M-line proteins specific to particular fiber types may interact differently with thick filaments to generate one or other type of lattice [8, 40, 41]. For example, M-protein, primarily responsible for the central stripe of the M-line (M1) and present only in fast (type IIB) fibers [42], could be the linker that defines relative thick filament orientation in a super lattice. When absent, as in slow and type IIA fibers, the thick filaments may default to a simple lattice arrangement. Alternatively, the lattice type could be dependent on the type of myosin present in the thick filament, which is known to vary with fiber type, as discussed earlier. In this case, the thick filament bridging proteins of the M-line may be the same in different fibers (e.g. myomesin is present in the M-lines of all fiber types [40, 41]), but interact differently with the thick filaments [43], depending on their myosin heavy chain type. Another thick filament structural protein with isoforms specific to different fiber types is myosin binding protein C (MyBP-C). Slow MyBP-C occurs in both fast (IIA and IIB) and slow (type I) fibers, while fast MyBP-C is present only in fast type IIB fibers (and absent from fast type IIA fibers) [11]. MyBP-C has been observed to bridge thick filaments to neighboring thin filaments [44], and thus can indirectly link thick filaments to their neighbors. The different forms of MyBP-C in different fiber types might link filaments differently, contributing to formation of the two types of lattice. If so, this would represent a novel function of MyBP-C, a protein whose role in skeletal muscle is not yet well understood [11, 45, 46]. MyBP-C is in fact long enough to directly link thick filaments, which could more directly influence lattice formation, although there is so far no evidence that such connections exist in the myofibril [44]. If this speculation is correct, disease mutations in skeletal MyBP-C and M-line proteins could affect phenotype at least in part through their putative role in defining filament lattice type.

## Acknowledgments

We thank Drs. John Woodhead, Peter Reiser and Elizabeth Ehler for discussions, and Michael Previs for sharing unpublished mass spectrometry data with us. This research was supported in part by NIH Grants AR072036 (to R.C.), AR067279 (to R.C. and D. Warshaw) and R01HL139883 (to Richard Moss and RC), and used resources of the Advanced Photon Source, a U.S. Department of Energy (DOE) Office of Science User Facility operated for the DOE Office of Science by Argonne National Laboratory under Contract No. DE-AC02-06CH11357. Use of the BioCAT Beamline 18ID was supported by grant P41 GM103622 from the National Institute of General Medical Sciences of the National Institutes of Health. Use of the Pilatus 3 1M detector was provided by grant 1S10OD018090-01 from NIGMS. We are grateful to the Core Electron Microscope Facility at UMass Medical School and its staff for use of their resources, and acknowledge funding from NIH grant 1S10ODO21580-01 for the ultramicrotome. The content of this work is solely the responsibility of the authors and does not necessarily reflect the official views of the National Institutes of Health.

## Supplementary Material

### Supplementary Results and Discussion

#### Mouse SOL and EDL

We wanted to test the generality of our conclusions concerning lattice/fiber type correlation, based on our rat observations, by studying the same question in mouse using X-ray diffraction and EM. The mouse is more difficult because its EDL and SOL have a greater mix of fiber types. While the EDL is ~100% fast (66% IIB, 21% IIDB, 8% IIAD [13]), the SOL is not a “pure” muscle, but a mixture of 60-70% fast and 30-40% slow [10]. Various studies have estimated fiber type amounts as: 37% type 1, 39% IIA, 19% IIAD [13]; 40% I, 50% IIA, 5% IIB/X [14]; and 34% I, 59% IIA, 6% IIB, 1% IIC [15].

If slow (type I) fibers have a simple lattice, and fast (type IIB/X/D) have a super lattice, as suggested by the rat results, this would predict a super lattice pattern for mouse EDL. However, the prediction for mouse SOL will depend on the lattice arrangement in type IIA fibers, as this is the predominant fiber type in mouse SOL; as discussed in the text, results from mouse diaphragm suggest that type IIA fibers have a simple lattice [33].

The X-ray results were suggestive, but not definitive. Mouse EDL showed little or no sampling on ML1, consistent with a super lattice that is very limited in extent and/or poorly ordered (Fig. S6B, E). Mouse SOL showed diffuse sampling on ML1 aligned with the 1,1 equatorial reflection (the strongest sampled reflection in the rat), while the rest of the layer line was unsampled. This is consistent with the presence of a simple lattice in a substantial fraction of fibers in mouse SOL. However, the diffuseness of the 1,1 sampling, and absence of sampling along the rest of the layer line, suggests that the simple lattice is not extensive/highly ordered and/or that some fibers in mouse SOL do not have a simple lattice. Lack of ordering seems more likely if we assume that type IIA fibers have a simple lattice. This possibility is supported by our observation that when mouse SOL was treated with blebbistatin, known to improve the ordering of myosin heads on the thick filaments, the sampling on the 1,1 row line became sharper, and a second spot appeared, on the 1,0 row line (the second strongest sampled reflection in rat) (Fig. S7). This suggests the presence of an (imperfectly ordered) simple lattice in most of the fibers in mouse SOL, supporting the view that both type I and IIA fibers have a simple lattice. This conclusion is also suggested by rat SOL, which, while containing up to 15% type IIA fibers [10, 11], shows a clear simple lattice, with no obvious sign of a super lattice. In contrast with the SOL, mouse EDL showed no change in sampling with blebbistatin. The relative weakness of simple lattice sampling in mouse SOL and absence of lattice sampling in EDL (Fig. S8) suggest that the lattices present in both muscles are not as ordered as in the rat.

We wanted to use electron microscopy of transverse sections of the bare zone to confirm the lattice arrangement present in different fiber types of the mouse, as we did for rat (Figs. 3, 4). We focused on the mouse SOL because of its mixture of fast and slow fibers. However, it is not possible to definitively identify fiber type in a mixed muscle based on transverse EM sections alone. One ultrastructural indicator of fiber type in transverse sections is the number of mitochondrial bundles lying immediately beneath the sarcolemma and in longitidudinal arrays parallel to the myofibrils [29, 47, 48]. These are common in slow type I and fast type IIA fibers, and absent from fast type IIB/D/X fibers. Using this feature as a guide, we attempted to distinguish type I/IIA from type IIB/D/X fibers in low magnification electron micrographs of mouse SOL. We then examined filament orientation at high magnification in bare zones of fibers identified in this way. Statistical analysis of relative filament orientations, similar to that carried out in Figs. 4 and S5, showed that “fast” fibers of the SOL (those with minimal mitochondrial bundles: IIB/D/X) had a mean deviation angle of 33.1 º when comparing filaments within a hexagon (Fig. S9A). As discussed for rat, this is consistent with a no-3-alike super lattice, in which two of every three adjacent filaments have the same angle and the third is rotated by 60º (Fig. 4C). However, it is also consistent with a random distribution of rotations. To distinguish these possibilities, we measured the relative orientations of pairs of adjacent filaments (cf. Fig. 4E, S5E). The results showed broad, similar height peaks for relative orientations close to 0 or 60º, supporting a no-3-alike super lattice in these fibers (Fig. S9B). If the fibers in EDL (essentially all fast) have the same lattice, why is super lattice sampling not seen in the X-ray diffraction pattern (Fig. S6)? Even in the best example of a super lattice (frog; [1]), the sampling is relatively weak (Fig. S4A, B), apparently due to the limited extent/disorder of the lattice [7, 8]. We suggest that the lattice in mouse may be more disordered than of rat or frog, and possibly also easily disrupted during specimen preparation.

We carried out a similar analysis on the mouse SOL fibers showing abundant mitochondria (assumed to be type I or IIA), where we expect a simple lattice. These showed a broad peak with an average deviation of 25.5º for filaments in a hexagon, higher than the 19.8º found for rat soleus (Fig. S9C) and the 0º expected for a perfect simple lattice. This was initially surprising, given the X-ray data for mouse SOL that suggested a (somewhat disordered) simple lattice (Figs. S6, 7). However, direct observation of relative orientations in these fibers showed many regions of similarly oriented triangles (Fig. S10), interspersed with others showing varying angles. This suggests that a simple lattice is indeed present, but that it is local and does not carry across an entire myofibril. We tested this idea by computing pairwise relative orientations as carried out for rat EDL and mouse SOL fast fibers. The results showed a substantially greater fraction of filaments with deviations close to 0º than close to 60º, clearly different from rat EDL and mouse SOL fast fibers, where the 0º and 60º had similar numbers. This supports the concept of local regions of simple lattice (Fig. S9D) rather than a random orientation of filaments, which would predict an even distribution of rotations from 0 to 60º. A disordered (i.e. local) simple lattice would be expected to give an X-ray diffraction pattern with sampling on the 1,0 and 1,1 row lines, but the sampling would be weaker/more diffuse than a for a perfect lattice, consistent with the observed weak sampling for the mouse SOL.

We conclude that overall, analysis of mouse SOL, by X-ray diffraction and EM, and mouse EDL by X-ray diffraction supports the general conclusion that low-fatigue fibers (type I and IIA) have a simple lattice while fatigable fibers (type IIB, D, X) have a super lattice. In no case is the lattice perfect: all have some degree of disorder (i.e. are limited in their extent).

---

**Figure S1.**
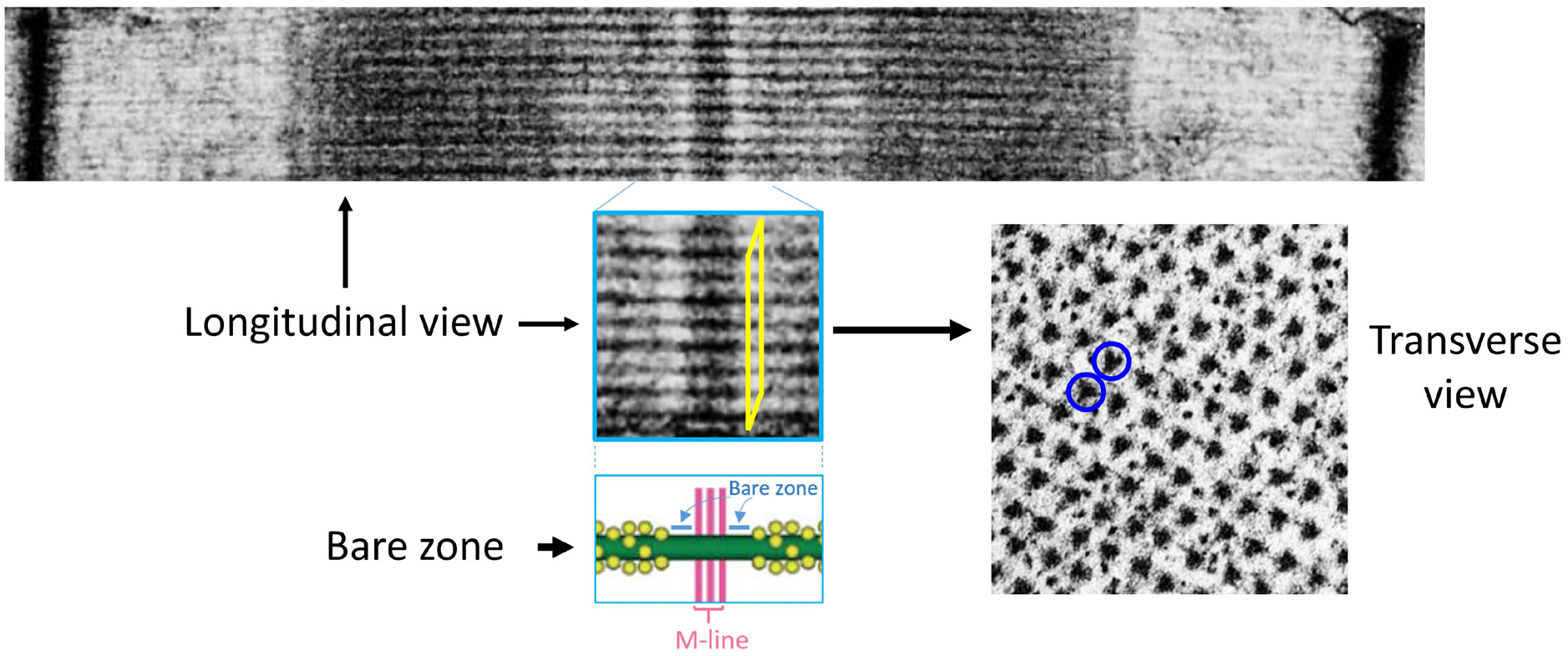
Transverse sectioning of the “bare zone”. Transverse sections of fixed and embedded muscle were cut and examined in myofibrils where the thick filaments showed triangular profiles (examples in blue circles). This occurs in the bare zones (yellow parallelogram), where M-line bridges and myosin heads are absent (cartoon).

**Figure S2.**
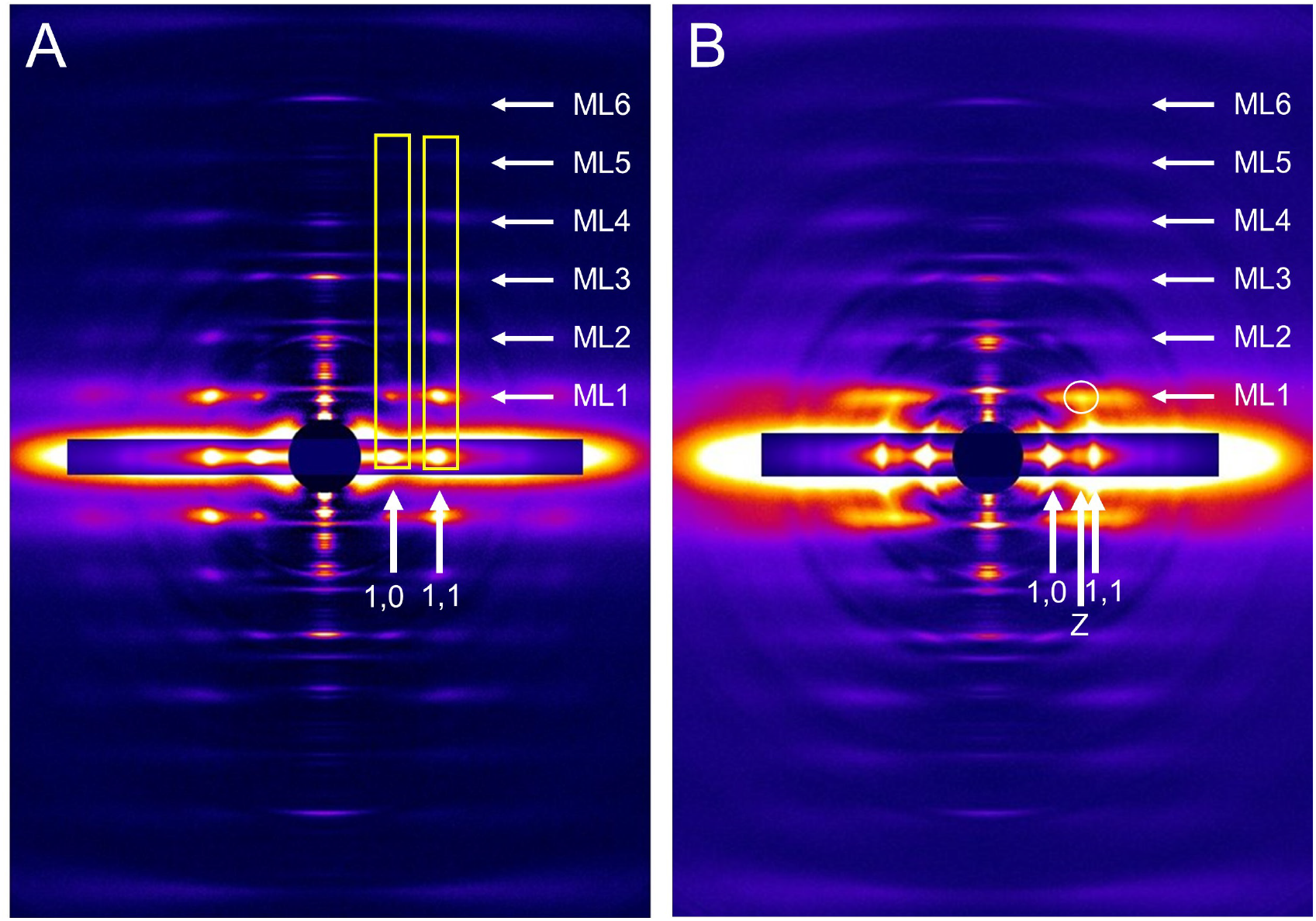
X-ray diffraction patterns of skinned rat SOL and EDL (compare with live patterns in Fig. 2). **A.** SOL – note clear lattice sampling on ML1 and higher layer lines at positions aligning with 1,0 and 1,1 equatorial reflections (cf. Fig. 2A). **B.** EDL – note much weaker lattice sampling and alignment of strongest (diffuse) spot on ML1 with Z-line reflection on equator (cf. Fig. 2B), consistent with super lattice. Note strong similarity to live patterns in Figs. 2, S3.

**Figure S3.**
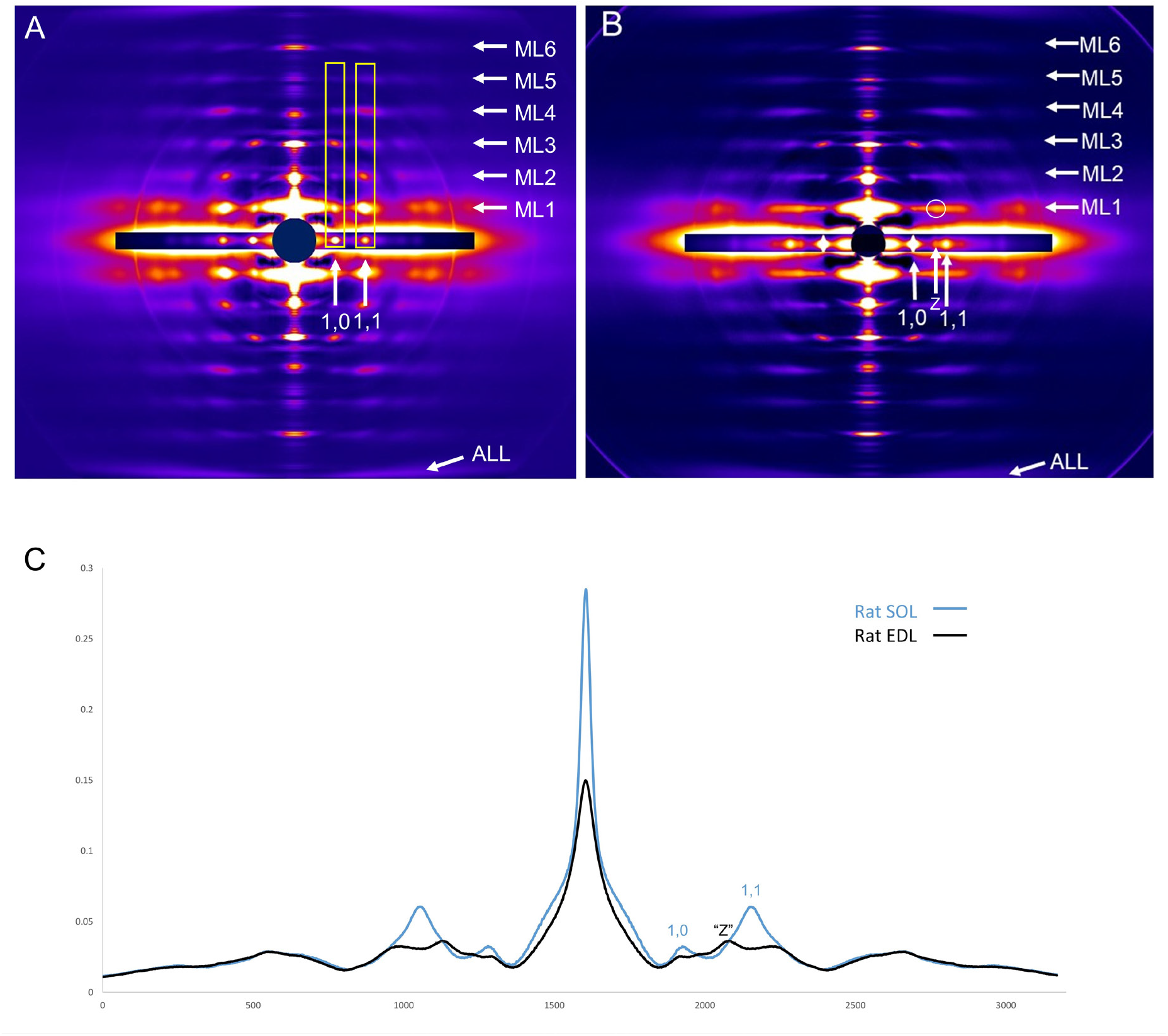
X-ray diffraction patterns of rat SOL (A) and EDL (B) (from Fig. 2) with intensity plot of first myosin layer line (C). ML1-6 indicate positions of myosin layer lines (repeat 43 nm), and ALL, actin 5.9 nm layer line. Horizontal inserts on equator show shorter exposures to reveal the high intensity equatorial 1,0 and 1,1 reflections. Prominent lattice sampling is seen in SOL at 1,0 and 1,1 radial positions (A, C), indicating a simple lattice of myosin filaments. Lattice sampling is weak in EDL, and in a different position from SOL, on the inner side of the 1,1 reflection (B, C), aligning approximately with the Z-line reflection on the equator (see Figs. 2, S4), consistent with a local super-lattice with disorder.

**Figure S4.**
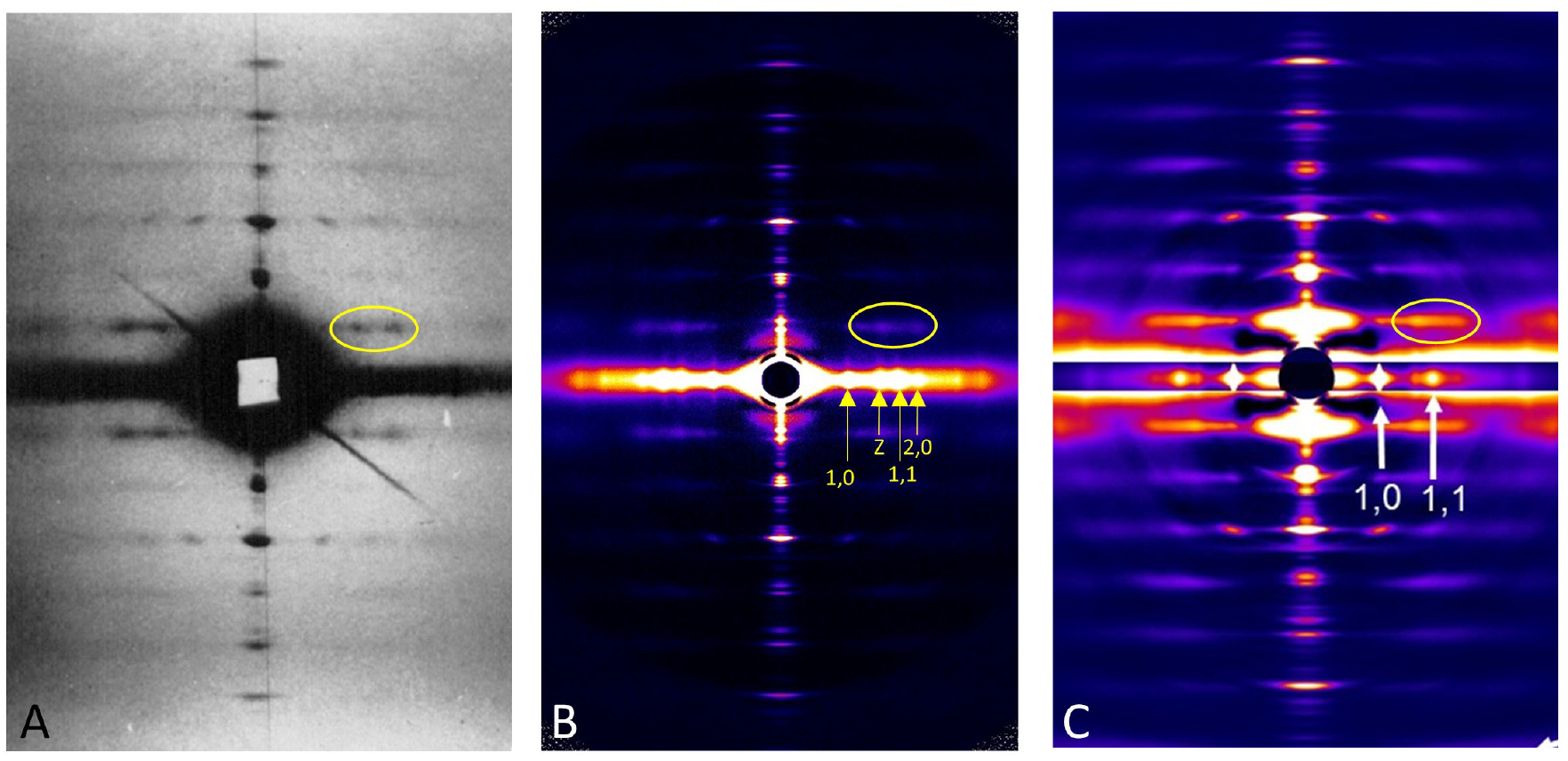
X-ray diffraction patterns demonstrating super lattice arrangement of thick filaments. **A.** Frog sartorius muscle (from Huxley and Brown [1], who first demonstrated the presence of a superlattice, with permission; ellipse indicates the two superlattice reflections first analyzed in this paper). **B.** Frog sartorius muscle. Pattern taken on same beamline as the rat muscles in this paper showing super lattice sampling as in (A) (T. Irving and H.E. Huxley, unpublished). **C.** Rat EDL (this study, from Fig. 2B) showing similar super lattice sampling in rat to that in frog.

**Figure S5.**
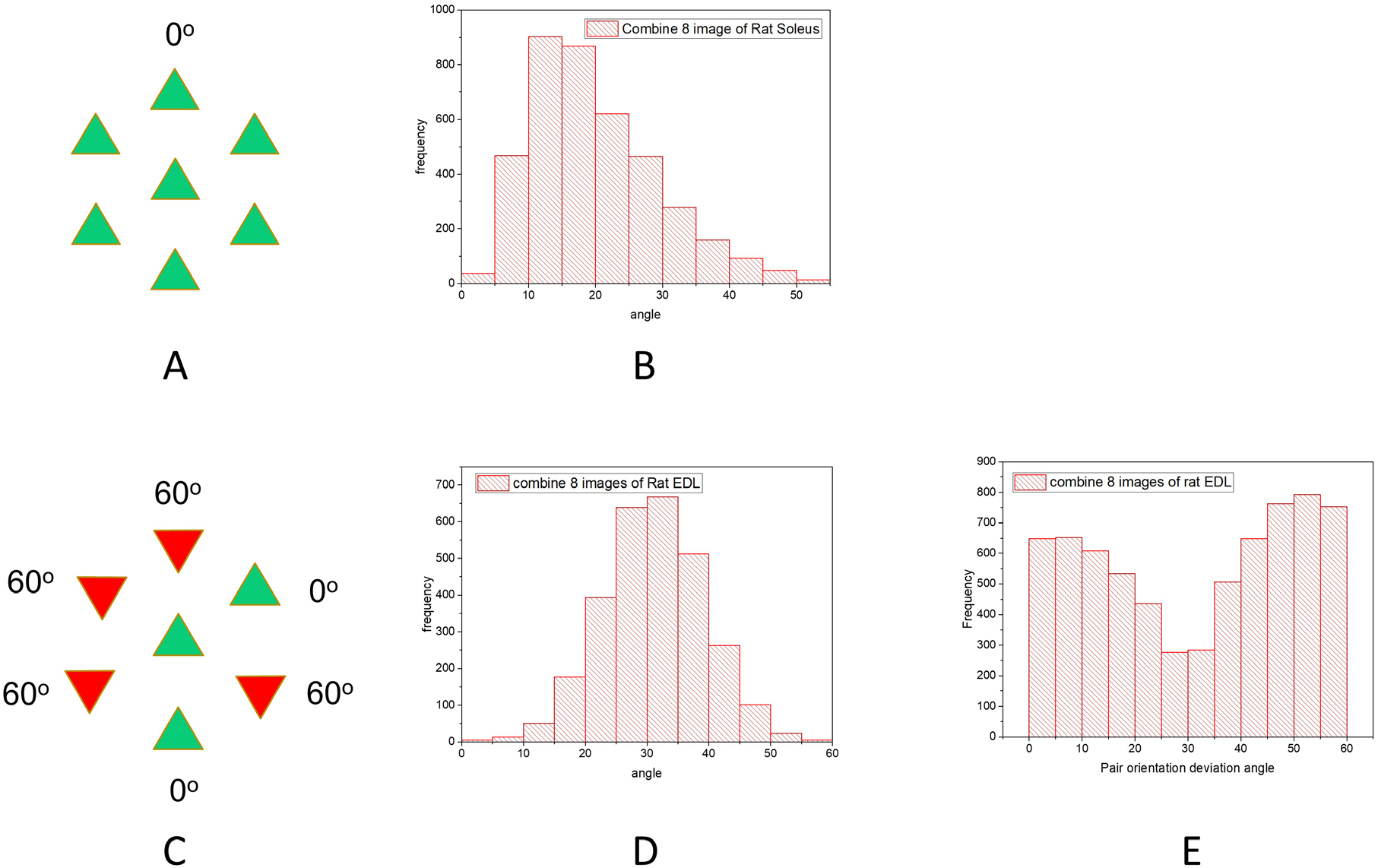
Analysis of filament orientations in rat SOL and EDL – all data (see Fig. 4). **A, C.** Filament orientations in simple lattice (**A**, all filaments have same orientation; mean deviation angle = 0º) and in no-3-alike super lattice (**C**: in any group of three filaments, in a line or forming a triangle, one filament is rotated by 60º with respect to the other two, which are identical; mean deviation angle for any three adjacent filaments = 30º). **B.** Histogram of deviation angles for all SOL micrographs analyzed. Average deviation angle = 19.8º, relatively close to the expected 0º for a simple lattice. The difference from 0º could be due to curvature of the lattice in many of the micrographs, which exaggerates the deviation angle (see Fig. 4 legend), to effects of muscle handling and EM processing, or it could reflect natural variation within the muscle. **D.** Deviation angle for three adjacent filaments in all EDL micrographs analyzed. The average deviation angle is 30.9º, similar to the expected 30º for a super lattice. **E.** Deviation angle for pairs of neighboring filaments in all EDL images analyzed. There are two peaks, close to 0 and 60º, as expected for a no-3-alike super lattice (see **C.**).

**Figure S6.**
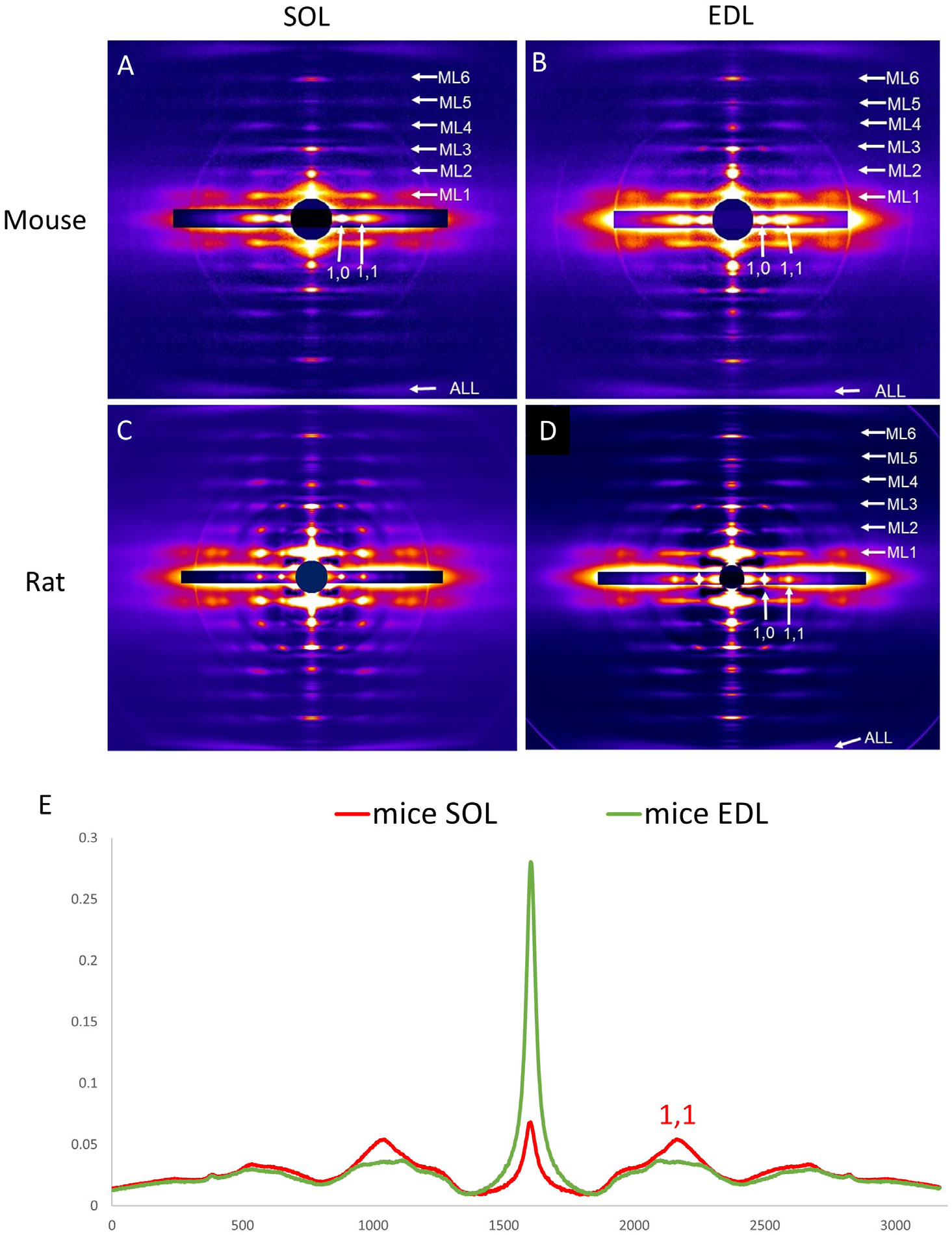
X-ray diffraction patterns of mouse SOL (A) and EDL (B) compared with rat SOL (C) and EDL (D) (from Fig. 2), with intensity plot of first myosin layer line in mouse (E). ML1-6 indicate positions of myosin layer lines (repeat 43 nm), and ALL, actin 5.9 nm layer line. 10 and 11 show positions of 10 and 11 equatorial reflections. Mouse EDL shows a relatively continuous density on ML1 **(B, E)**, while mouse SOL shows a mixture of continuous density with significant, slightly diffuse sampling at the 1,1 radial position **(A, E)**. Sampling is stronger in the rat than the mouse (see Fig. S7).

**Figure S7.**
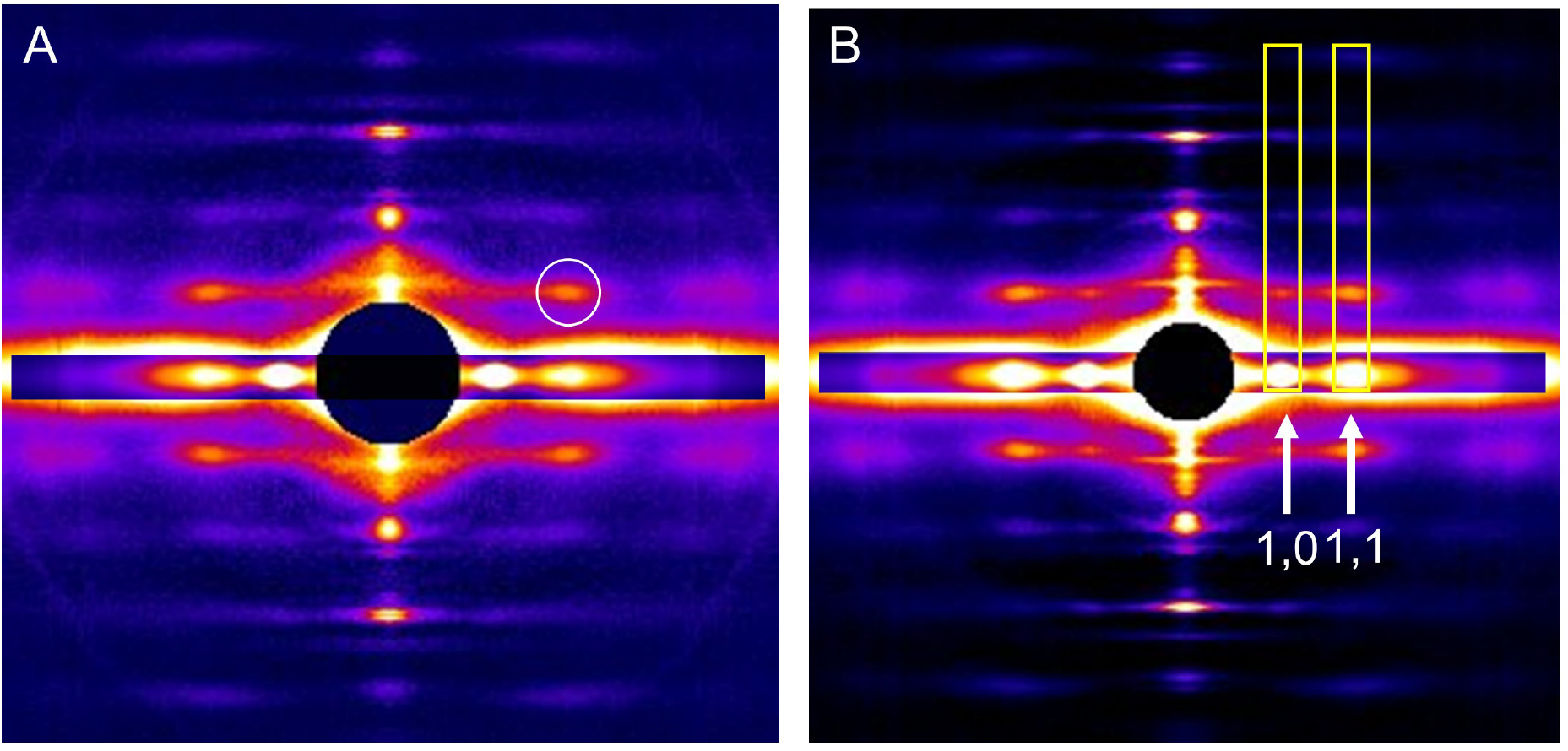
X-ray diffraction of intact mouse SOL. In **(A)**, lattice sampling on ML1 (circle) was aligned with the 1,1 equatorial reflection, but there was no sign of 1,0 sampling. When the muscle was treated with blebbistatin **(B)**, this lattice sampling became stronger and sharper, and a lattice spot on ML1 was now seen aligned with the 1,0 equatorial reflection (rectangles). The enhanced lattice sampling is presumably due to improved helical ordering of the myosin heads which sharpens myosin layer lines [2].

**Figure S8.**
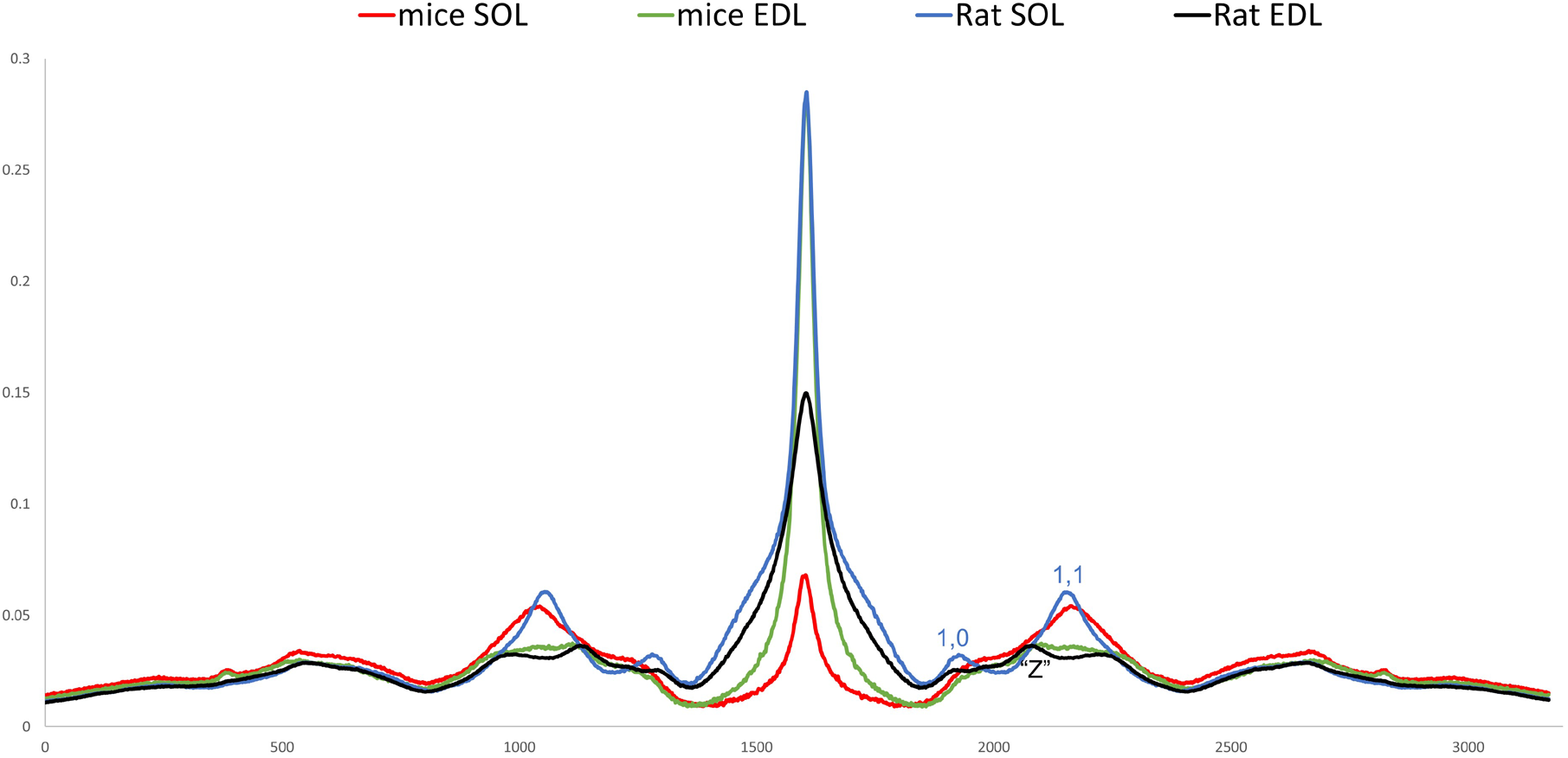
Comparison of intensity plots of ML1 in rat and mouse SOL and EDL. Both SOL muscles show a peak in line with the 1,1 equatorial reflection. The peak with rat is sharper than with mouse. Rat also shows a peak at the 1,0 radial position. Sampling with the EDL muscles is weaker. However, the rat shows a small but clear peak in line with the Z-line equatorial reflection, on the inner side of the 1,1 and another peak on the outer side of the 1,1. These reflections are characteristic of a super lattice (see Fig. S4).

**Figure S9.**
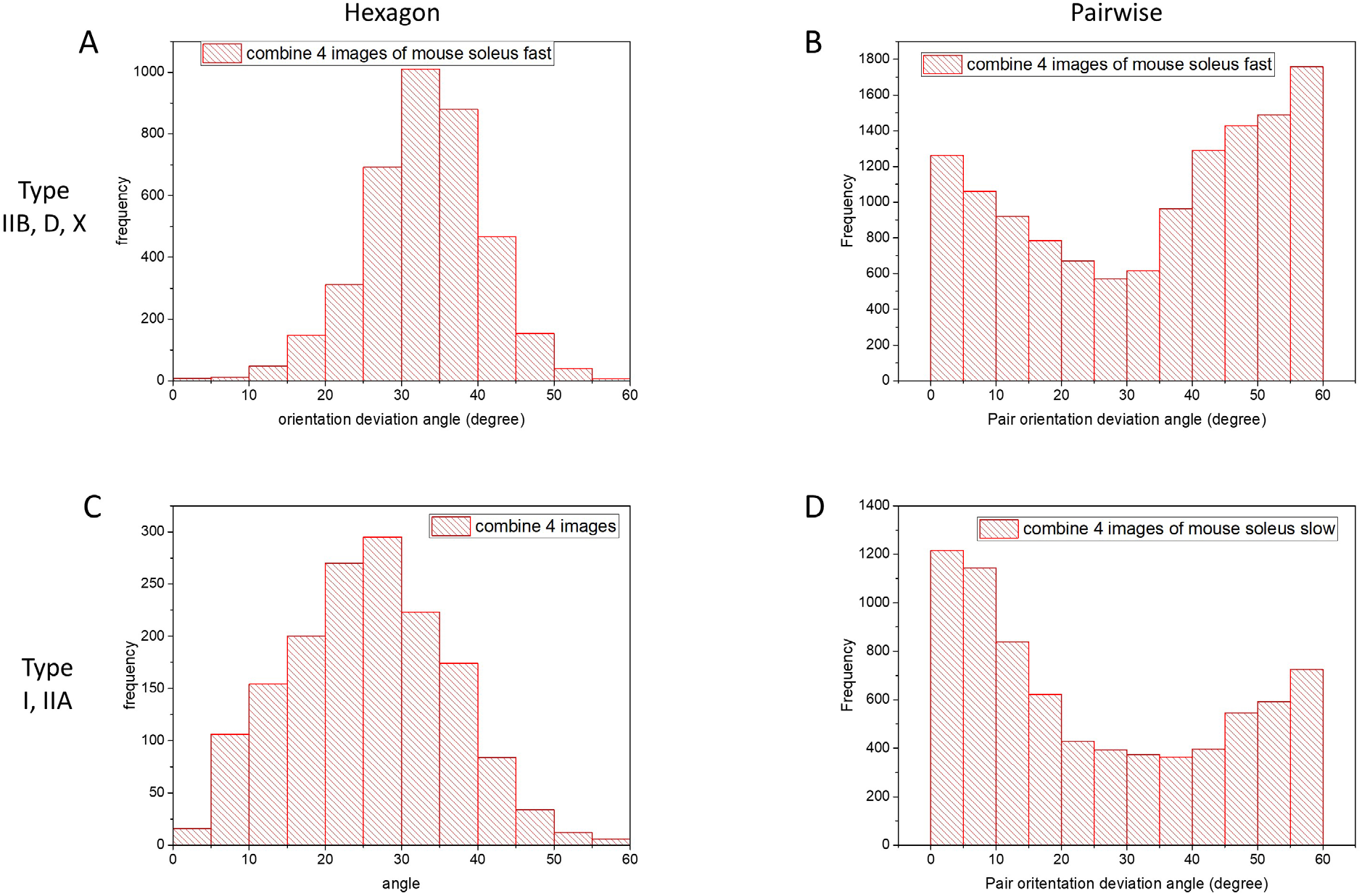
Average angular deviation of mouse SOL thick filaments in fast (type IIB, D, X) and slow (type I) and type fast (type IIA) fibers. **(A)** and **(C)** show average deviation for filaments in hexagons centered on the reference filament (cf. Fig. S5B, D). **(B)** and **(D**) show pairwise comparison of filaments. Note that **(A)** and **(C)** have average deviation angles of 30.9 and 25.5º respectively. For the pairwise comparison, **(B)** has similar height peaks close to 0 and 60º, as expected for a no-3-alike super lattice, while **(D)** shows a substantially higher peak close to 0º than that at 60º, more consistent with a limited extent/disordered simple lattice (see Fig. S10).

**Figure S10.**
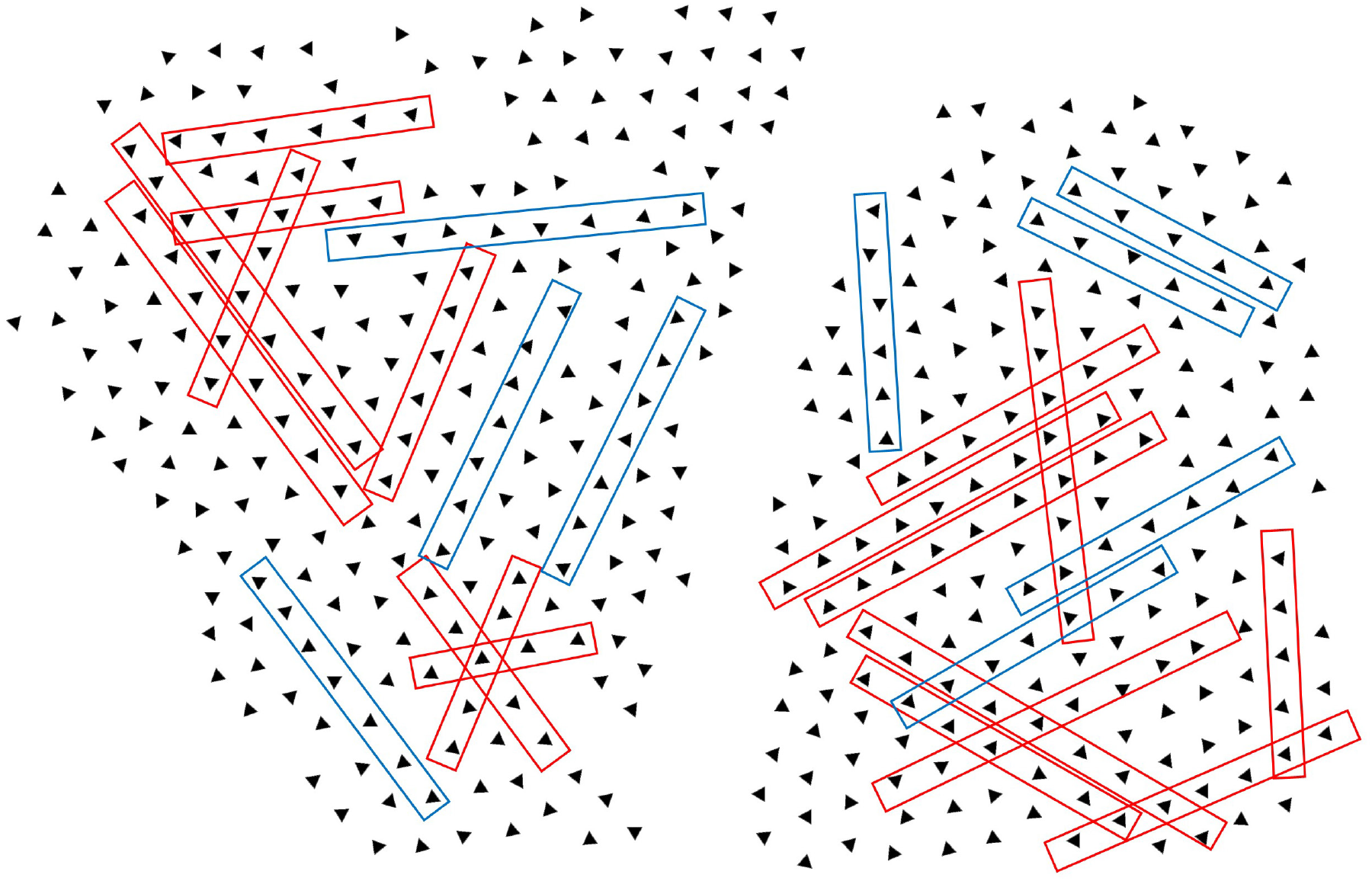
Thick filament orientations in type I/IIA fibers of mouse SOL. Fiber type was tentatively identified as type I/IIA based on high mitochondrial content. Thick filament orientations in two myofibrils were determined using the same methods as for Figs. 3D, F. Rectangles show local regions where filaments have similar orientations (simple lattice; red) interspersed with varying orientations (blue).

## References

1. Huxley, H.E., and Brown, W. (1967). The low-angle x-ray diagram of vertebrate striated muscle and its behaviour during contraction and rigor. J. Mol. Biol 30, 383–434.

2. Xu, S., White, H.D., Offer, G.W., and Yu, L.C. (2009). Stabilization of helical order in the thick filaments by blebbistatin: further evidence of coexisting multiple conformations of myosin. Biophys J 96, 3673–3681.

3. Luther, P.K., Squire, J.M., and Forey, P.L. (1996). Evolution of myosin filament arrangements in vertebrate skeletal muscle. J. Morphol 229, 325–335.

4. Harford, J., and Squire, J. (1986). "Crystalline" myosin cross-bridge array in relaxed bony fish muscle. Biophys. J 50, 145–155.

5. Huxley, H.E. (1968). Structural difference between resting and rigor muscle; evidence from intensity changes in the lowangle equatorial x-ray diagram. J. Mol. Biol 37, 507–520.

6. Steven, A.C., Baumeister, W., Johnson, L.N., Perham, R.N. (2016). Molecular Biology of Assemblies and Machines, (New York and London: Garland Science).

7. Luther, P.K., and Squire, J.M. (1980). Three-dimensional structure of the vertebrate muscle A-band. II. The myosin filament superlattice. J. Mol. Biol 141, 409–439.

8. Luther, P.K., and Squire, J.M. (2014). The intriguing dual lattices of the Myosin filaments in vertebrate striated muscles: evolution and advantage. Biology (Basel) 3, 846–865.

9. Luther, P.K., Munro, P.M., and Squire, J.M. (1981). Three-dimensional structure of the vertebrate muscle A-band. III. M-region structure and myosin filament symmetry. J. Mol. Biol 151, 703–730.

10. Wigston, D.J., and English, A.W. (1992). Fiber-type proportions in mammalian soleus muscle during postnatal development. J. Neurobiol 23, 61–70.

11. Li, A., Nelson, S., Rahmanseresht, S., Braet, F., Cornachione, A.S., Previs, S.B., O’Leary, T.S., McNamara, J.W., Rassier, D.E., Sadayappan, S., et al. (2019). Skeletal MyBP-C isoforms tune the molecular contractility of divergent skeletal muscle systems. BioRxiv https://www.biorxiv.org/content/10.1101/676601v1.

12. Eng, C.M., Smallwood, L.H., Rainiero, M.P., Lahey, M., Ward, S.R., and Lieber, R.L. (2008). Scaling of muscle architecture and fiber types in the rat hindlimb. J. Exp. Biol 211, 2336–2345.

13. Augusto, V., Padovani, C.R., and Campos, G.E.R. (2004). Skeletal muscle fiber types in C57BL6J mice. Braz. J. Morphol. Sci 21, 89–94.

14. Luedeke, J.D., McCall, R.D., Dillaman, R.M., and Kinsey, S.T. (2004). Properties of slow- and fast-twitch skeletal muscle from mice with an inherited capacity for hypoxic exercise. Comp Biochem PhysiolA Mol Integr Physiol 138, 373–382.

15. Totsuka, Y., Nagao, Y., Horii, T., Yonekawa, H., Imai, H., Hatta, H., Izaike, Y., Tokunaga, T., and Atomi, Y. (2003). Physical performance and soleus muscle fiber composition in wild-derived and laboratory inbred mouse strains. J Appl Physiol (1985. 95, 720–727.

16. Fischetti, R., Stepanov, S., Rosenbaum, G., Barrea, R., Black, E., Gore, D., Heurich, R., Kondrashkina, E., Kropf, A.J., Wang, S., et al. (2004). The BioCAT undulator beamline 18ID: a facility for biological non-crystalline diffraction and X-ray absorption spectroscopy at the Advanced Photon Source. J. Synchrotron. Radiat 11, 399–405.

17. Jiratrakanvong, J., Shao, J., Menendez, M., Li, X., Li, J., Ma, W., Agam, G., and Irving, T. (2018). MuscleX: software suite for diffraction X-ray imaging V1.13.1.

18. Frank, J. (2006). Three-dimensional electron microscopy of macromolecular assemblies: visualization of biological molecules in their native state, 2nd edition Edition, (New York: Oxford University Press).

19. Scheres, S.H. (2012). RELION: implementation of a Bayesian approach to cryo-EM structure determination. J. Struct. Biol 180, 519–530.

20. Frank, J., Radermacher, M., Penczek, P., Zhu, J., Li, Y., Ladjadj, M., and Leith, A. (1996). SPIDER and WEB: processing and visualization of images in 3D electron microscopy and related fields. J Struct Biol 116, 190–199.

21. Squire, J.M., Roessle, M., and Knupp, C. (2004). New X-ray diffraction observations on vertebrate muscle: organisation of C-protein (MyBP-C) and troponin and evidence for unknown structures in the vertebrate A-band. J. Mol. Biol 343, 1345–1363.

22. Reconditi, M. (2006). RECENT IMPROVEMENTS IN SMALL ANGLE X-RAY DIFFRACTION FOR THE STUDY OF MUSCLE PHYSIOLOGY. Rep. Prog. Phys 69, 2709–2759.

23. Linari, M., Brunello, E., Reconditi, M., Fusi, L., Caremani, M., Narayanan, T., Piazzesi, G., Lombardi, V., and Irving, M. (2015). Force generation by skeletal muscle is controlled by mechanosensing in myosin filaments. Nature 528, 276–279.

24. Xu, S., Offer, G., Gu, J., White, H.D., and Yu, L.C. (2003). Temperature and ligand dependence of conformation and helical order in myosin filaments. Biochemistry 42, 390–401.

25. Hamalainen, N., and Pette, D. (1993). The histochemical profiles of fast fiber types IIB, IID, and IIA in skeletal muscles of mouse, rat, and rabbit. J Histochem Cytochem 41, 733–743.

26. Salviati, G., Betto, R., and Danieli Betto, D. (1982). Polymorphism of myofibrillar proteins of rabbit skeletal-muscle fibres. An electrophoretic study of single fibres. Biochem J 207, 261–272.

27. Lutz, G.J., Bremner, S., Lajevardi, N., Lieber, R.L., and Rome, L.C. (1998). Quantitative analysis of muscle fibre type and myosin heavy chain distribution in the frog hindlimb: implications for locomotory design. J Muscle Res Cell Motil 19, 717–731.

28. Gutmann, E. (1966). "Slow" and "Fast" Muscle Fibers. MCV Quarterly 2, 78–81.

29. Schiaffino, S., and Reggiani, C. (2011). Fiber types in mammalian skeletal muscles. Physiol Rev 91, 1447–1531.

30. Yamaguchi, M., Takemori, S., Kimura, M., Nakahara, N., Ohno, T., Yamazawa, T., Yokomizo, S., Akiyama, N., and Yagi, N. (2016). Approaches to physical fitness and sports medicine through X-ray diffraction analysis of striated muscle. J Phys Fitness Sports Med 5, 47–55.

31. Horiuti, K., Yagi, N., and Takemori, S. (1997). Mechanical study of rat soleus muscle using caged ATP and X-ray diffraction: high ADP affinity of slow cross-bridges. J. Physiol (Lond) 502 (Pt 2), 433–447.

32. Matsubara, I., Yagi, N., Saeki, Y., and Kurihara, S. (1991). Cross-bridge movement in fast and slow skeletal muscles of the chick. J. Physiol (Lond) 441, 113–120.

33. Iwamoto, H., Wakayama, J., Fujisawa, T., and Yagi, N. (2003). Static and dynamic x-ray diffraction recordings from living mammalian and amphibian skeletal muscles. Biophys. J. 85, 2492–2506.

34. Coirault, C., Chemla, D., Pery-Man, N., Suard, I., and Lecarpentier, Y. (1995). Effects of fatigue on force-velocity relation of diaphragm. Energetic implications. Am J Respir Crit Care Med 151, 123–128.

35. Soukup, T., Zacharova, G., and Smerdu, V. (2002). Fibre type composition of soleus and extensor digitorum longus muscles in normal female inbred Lewis rats. Acta Histochem 104, 399–405.

36. Luther, P.K., Munro, P.M.G., and Squire, J.M. (1995). Muscle ultrastructure in the teleost fish. Micron 26, 431–459.

37. Polla, B., D’Antona, G., Bottinelli, R., and Reggiani, C. (2004). Respiratory muscle fibres: specialisation and plasticity. Thorax 59, 808–817.

38. Rome, L.C., Funke, R.P., Alexander, R.M., Lutz, G., Aldridge, H., Scott, F., and Freadman, M. (1988). Why animals have different muscle fibre types. Nature 335, 824–827.

39. Squire, J.M., Bekyarova, T., Farman, G., Gore, D., Rajkumar, G., Knupp, C., Lucaveche, C., Reedy, M.C., Reedy, M.K., and Irving, T.C. (2006). The myosin filament superlattice in the flight muscles of flies: A-band lattice optimisation for stretch-activation? J Mol Biol 361, 823–838.

40. Lange, S., Pinotsis, N., Agarkova, I., and Ehler, E. (2019). The M-band: The underestimated part of the sarcomere. Biochim Biophys Acta Mol Cell Res.

41. Schoenauer, R., Lange, S., Hirschy, A., Ehler, E., Perriard, J.C., and Agarkova, I. (2008). Myomesin 3, a novel structural component of the M-band in striated muscle. J Mol Bio 376, 338–351.

42. Obermann, W.M., Gautel, M., Steiner, F., van der Ven, P.F., Weber, K., and Furst, D.O. (1996). The structure of the sarcomeric M band: localization of defined domains of myomesin, M-protein, and the 250-kD carboxy-terminal region of titin by immunoelectron microscopy. J. Cell Biol 134, 1441–1453.

43. Pask, H.T., Jones, K.L., Luther, P.K., and Squire, J.M. (1994). M-band structure, M-bridge interactions and contraction speed in vertebrate cardiac muscles. J. Muscle Res. Cell Motil 15, 633–645.

44. Luther, P.K., Winkler, H., Taylor, K., Zoghbi, M.E., Craig, R., Padron, R., Squire, J.M., and Liu, J. (2011). Direct visualization of myosin-binding protein C bridging myosin and actin filaments in intact muscle. Proc Natl Acad Sci U S A 108, 11423–11428.

45. Geist, J., Ward, C.W., and Kontrogianni-Konstantopoulos, A. (2018). Structure before function: myosin binding protein-C slow is a structural protein with regulatory properties. FASEB J, fj201800624R.

46. Robinett, J.C., Hanft, L.M., Geist, J., Kontrogianni-Konstantopoulos, A., and McDonald, K.S. (2019). Regulation of myofilament force and loaded shortening by skeletal myosin binding protein C. J Gen Physiol 151, 645–659.

47. Schiaffino, S., Hanzlikova, V., and Pierobon, S. (1970). Relations between structure and function in rat skeletal muscle fibers. J Cell Biol 47, 107–119.

48. Gauthier, G.F. (1986). Skeletal muscle fiber types. In Myology, A.G.B. Engel, B.Q., ed. (New York: McGraw-Hill Book Company), pp. 255–283.

